# What makes eye movements effective retrieval cue: The contribution of visual and the motor components to memory performance

**DOI:** 10.1101/2022.11.26.517864

**Authors:** Keren Taub, Yonatan Goshen-Gottstein, Shlomit Yuval-Greenberg

## Abstract

During memory retrieval, people tend to reenact the same eye movements performed when memorized items were first displayed and to gaze at similar locations. This was hypothesized to reflect the role of eye movements as retrieval cues. However, it is unknown what is it about eye movements that makes them effective retrieval cues. Here, we examine, for the first time, the individual and combined contributions of the visual and the motor components of eye movements to memory retrieval. Results (N=70) revealed a non-additive benefit of both components of eye movements to memory performance. Additionally, we found that individuals who gained from one component, were more likely to gain from the other as well. Together, these findings unravel the central role of eye movements in episodic memory; they show how the visual and motor components are integrated into a single effective memory retrieval cue and how this integration varies among individuals.

## Introduction

It is a wide consensus among researchers that the main purpose of eye movements is to guide the center of vision, the fovea, towards locations of interest. Yet studies have frequently shown that while performing various non-visual tasks, participants often shift their eyes to empty locations where relevant information may have previously appeared, but no longer does^1–5^. Specifically, this “looking-at-nothing” phenomenon was demonstrated in free-recall episodic memory tasks. A highly consistent finding is that at the time of retrieval of episodic memories, people tend to shift their gaze to the same locations where they had fixated during encoding, even when those locations are empty at the time of retrieval^2,6–12^. In addition, when participants were asked to verbally describe a scene from memory or to form mental images of visual stimuli that they have previously seen, they tend to recreate the same eye movements pattern that they had performed while initially viewing the images^10,13–15^. Similar findings were obtained when participants were asked to recall visual memories ^13,14,16^.

These behaviors – of performing gaze shifts to locations where memorized items were first presented and reenacting the eye movements that were performed while viewing these times - were found to be associated with enhancement in the ability to remember these items. For example, recognition judgements were found to be faster when items were retrieved while performing horizontal eye movements in a similar direction (left/right) as in encoding, compared to when different eye movements were performed^17^. A study by Johansson and Johansson (2014) provided further support for this claim by showing that participants were more likely to correctly answer questions regarding a target object when they were instructed to gaze at the location where the object has been previously presented, relative to when they were instructed to gaze at a different location. Another study showed that reenacting the same gaze path during encoding and retrieval improves recall of simple checkerboard patterns, compared to when no reenactment was performed^18^. Furthermore, it was found that recreating the same gaze locations enhances memory performance even when the retrieved information is not visual^19^. This eye movement-related memory-enhancement was explained via the influential *encoding specificity principle*. This principle states that as similarity between encoding and retrieval conditions increases, memory performance improves^20,21^. According to this principle, when a retrieval cue is effective in reinstating the processes or actions undertaken at encoding, encoded memories resurface. The finding that eye movements performed during retrieval are similar to those performed during encoding, and the finding that this effect enhances memory were thus hypothesized to indicate that eye movements play a role as memory retrieval cues^16,18,22^.

Despite abundant evidence on the key role of eye movements in memory, relatively little is known about the mechanism by which eye movements function as a retrieval cue. One theory suggests that the eyes are shifted to locations where memorized items were previously presented, as a result of the activation of memory representations of the spatial indices of those items^23,24^. According to this theory, the mentioning of a previously seen object activates a set of memorized characteristics of that object, including its spatial location. The eyes are shifted towards that location with the goal of gathering more information, but no relevant information is collected because the object is no longer there^12^. This theory view eye movements as the consequence of memory activation rather than a retrieval cue. It does not explain what causes the improvement in memory performance when people are looking at empty locations where items were previously presented but no longer do: Is it the visual input that accompanies the eye movements, or the motor action that produced them? More specifically, with each eye movement, the visual input on the retina refreshes— leading to new visual stimulation (the visual component of eye movements). Although the memorized object is no longer present during retrieval, the rest of the visual scene – the screen and its surroundings – still is. This visual information, when matched between encoding and retrieval, could play a role in memory retrieval. In addition, the extraocular muscles contract in a unique pattern of contraction (the motor component of eye movements). It could be hypothesized that when a similar motor action is produced during encoding and retrieval, this match could also contribute to memory retrieval. Both of these components of eye movements could influence memory retrieval, separately or together, and their relative influence could vary between individuals. The goal of this study was to examine, for the first time, the role of eye movements in memory retrieval while disentangling their visual and motor components and investigate how they manifest in different conditions and different individuals. The findings of this study explain the link between eye movements and memory retrieval and unravel what makes eye movements effective retrieval cues: is it the visual input that accompanies them, or the motor action that produces them?

## Results

In an experiment with 70 participants, we investigated the separate and combined contributions of the visual and the motor components of eye movements to memory performance, and examined individual differences in the behavioral gain from these components. In each trial of an encoding phase, participants performed two sequential horizontal saccades, and were then presented with a target word at the final landing position. Trials varied both in the direction of the saccades (left/right), and in their vertical location on to the screen (top/bottom of the screen). During a following retrieval phase, participants performed the same eye movements task, but they were also asked to indicate whether the presented word (*lure*) had been presented before during encoding (recognition task). A match between encoding and retrieval in the direction of saccades (either left or right), which resulted in the performance of the same eye movement prior to the same target, was considered a motor cue. A match between encoding and retrieval in the vertical location of the presentation (top or bottom), which resulted in the same visual input at the presentation of the same target, was considered a visual cue. Each retrieval trial matched its accompanying encoding trial in the direction of the saccades (motor-only cue; M), the vertical location (visual-only cue; V), both (visual-motor cue; VM) or neither (no cue). Trial procedure is depicted in Figure 1.

**Figure 1.**
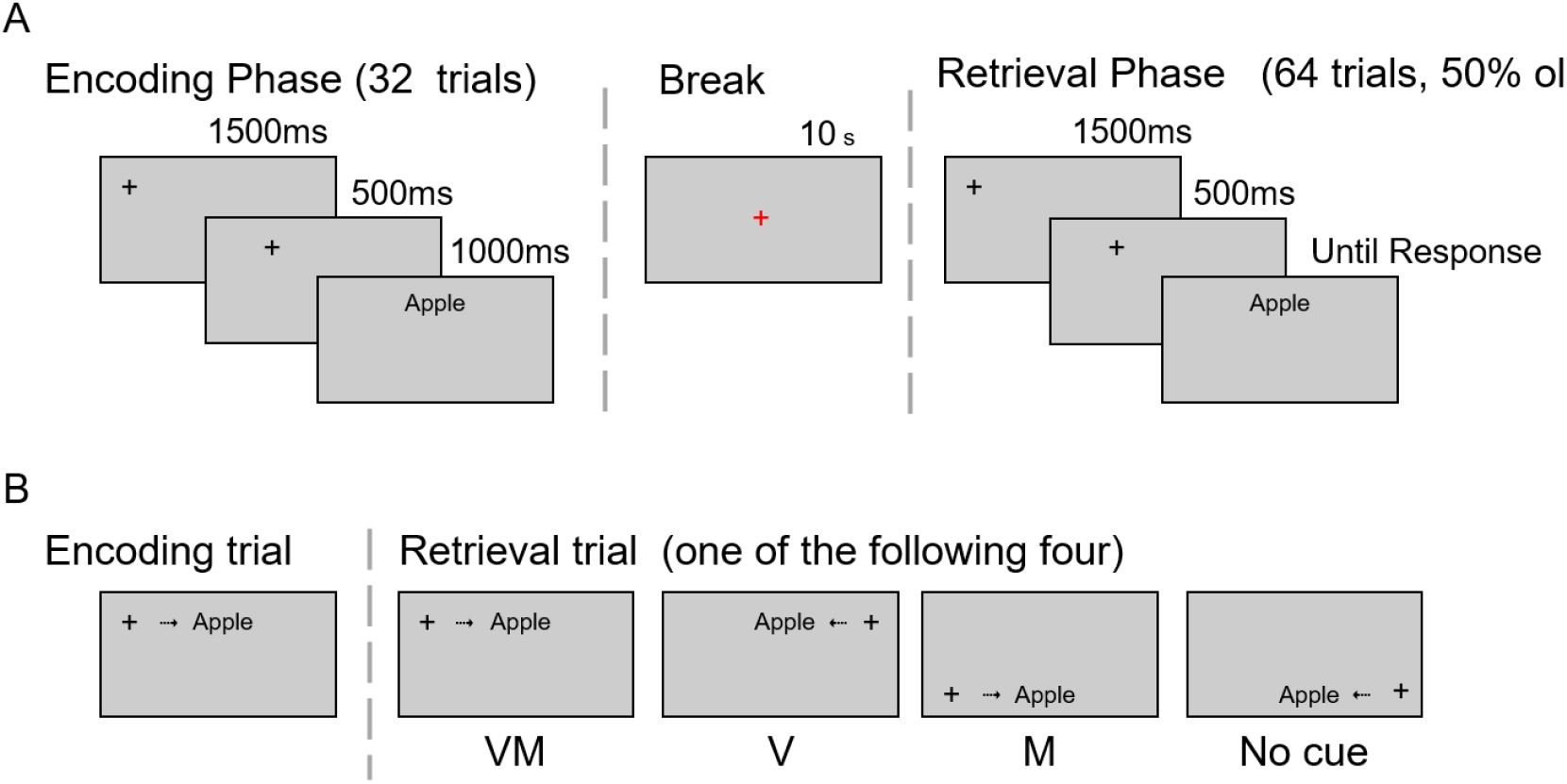
Experimental procedure. A) Each block included an encoding phase, a break (32 trials, followed by a short break and then a retrieval phase (64 trials, 50% of them presented in the encoding phase). B) A sample trial. The condition was determined according to the type of match between the corresponding encoding and retrieval trials: A match in both vertical position of the presentation and the direction of saccades provided both motor and visual cues (VM); a match in the vertical position of the presentation but not in the direction of saccades provided a visual cue (V); a match in the direction of saccades but not the vertical position of the presentation provided a motor cue (M); incongruity in both the vertical location and the direction of saccades provided no cue (No cue).

### The contribution of eye movements related cues

To test for the general contribution of cues related to eye movements, we compared recognition performance on the three cues-conditions (VM, V, M) to the No-cues condition. Memory performance was analyzed based signal-detection theory (SDT)^25^, in order to control for personal biases. We used the d_a_ index, which is similar to the known d’ index but does not rely on the assumption equal variance of the lure and target distributions^26^ (for a full explanation, see methods section).

Results showed that, as predicted, the presence of cues related to eye movements (visual and motor) were beneficial to memory performance relative to no cue (independent planned contrasts; 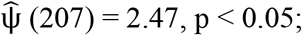 **Fig. 2**). Next, we tested for the beneficial effect of the combination of the two cues as compared to a single cue. To this end, we compared the VM condition with the two single-cue conditions (V, M) collapsed. Results indicated that there was no significant difference in memory performance when both cues were present compared to the presentation of a single cue condition (planned contrast; 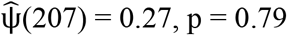), and this null effect was supported by Bayesian analysis (Bayesian paired samples of VM compared with the average of V and M; BF_01_ = 7.33). There was also no evidence for difference in performance between trials of the visual and the motor single cues (V and M) 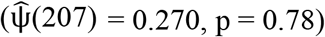. This null effect too was supported by Bayesian analysis (Bayesian paired samples comparing V and M; BF_01_ = 7.36). This suggests that, while the existence of cues benefited performance, a single cue, either motor or visual, was sufficient to achieve this benefit to its full extent.

**Figure 2.**
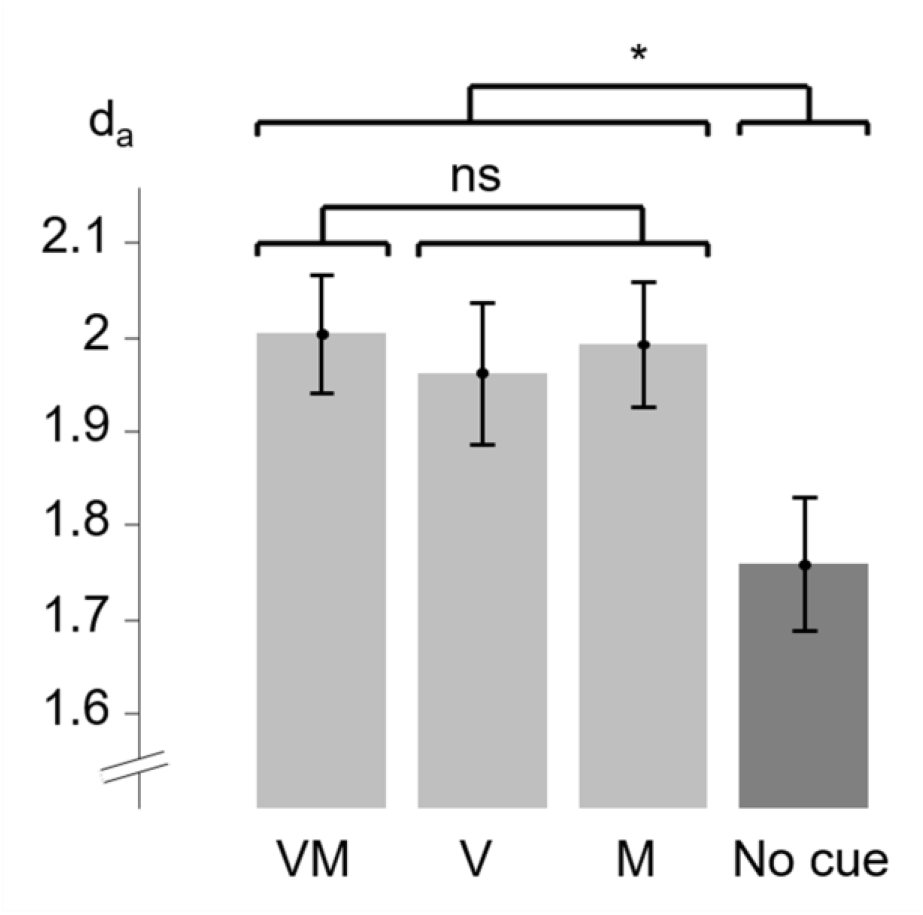
Memory performance. The da values of the four conditions: visual-motor cues (VM), visual cue (V), motor cue (M) and no cue. Error bars designate ±1 standard error from the condition’s mean, corrected for within-subject variability (Cousineau & O’Brien, 2014). * p < .05.

### Individual differences in the contribution of single cues

Our next aim was to examine differences among participants in the way the used eye movements as retrieval cues and how much gained from each of the cue types (visual or motor). To isolate the unique gain for performance of each single cue, while controlling for the participant’s general memory ability, we calculated for each participant and cue-condition (visual and motor) the difference in the d_a_ values between each of the single cue conditions and the No-cues condition (Equation (1) and Equation (1)).

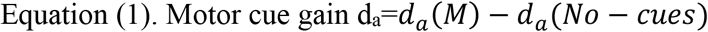

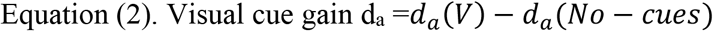

We then computed the Pearson correlation between the motor-cue gain and the visual-cue gain. We found a large positive correlation between gain of the visual cue and gain of the motor cue (Pearson r = 0.55, p < 0.001; **Fig. 3**). This suggests that participants who benefited from one cue when presented alone, tended do benefit to a similar extent also from the other cue when presented alone. Surprisingly, these findings also reveal that participants who had a negative gain for one cue, that is, performed worse with this cue than without it, tended to likewise show a negative gain for the other cue. This suggests that while some participants were able to gain from either cue, some performed worse in the presence of these cues compared with the No-cues condition.

**Figure 3.**
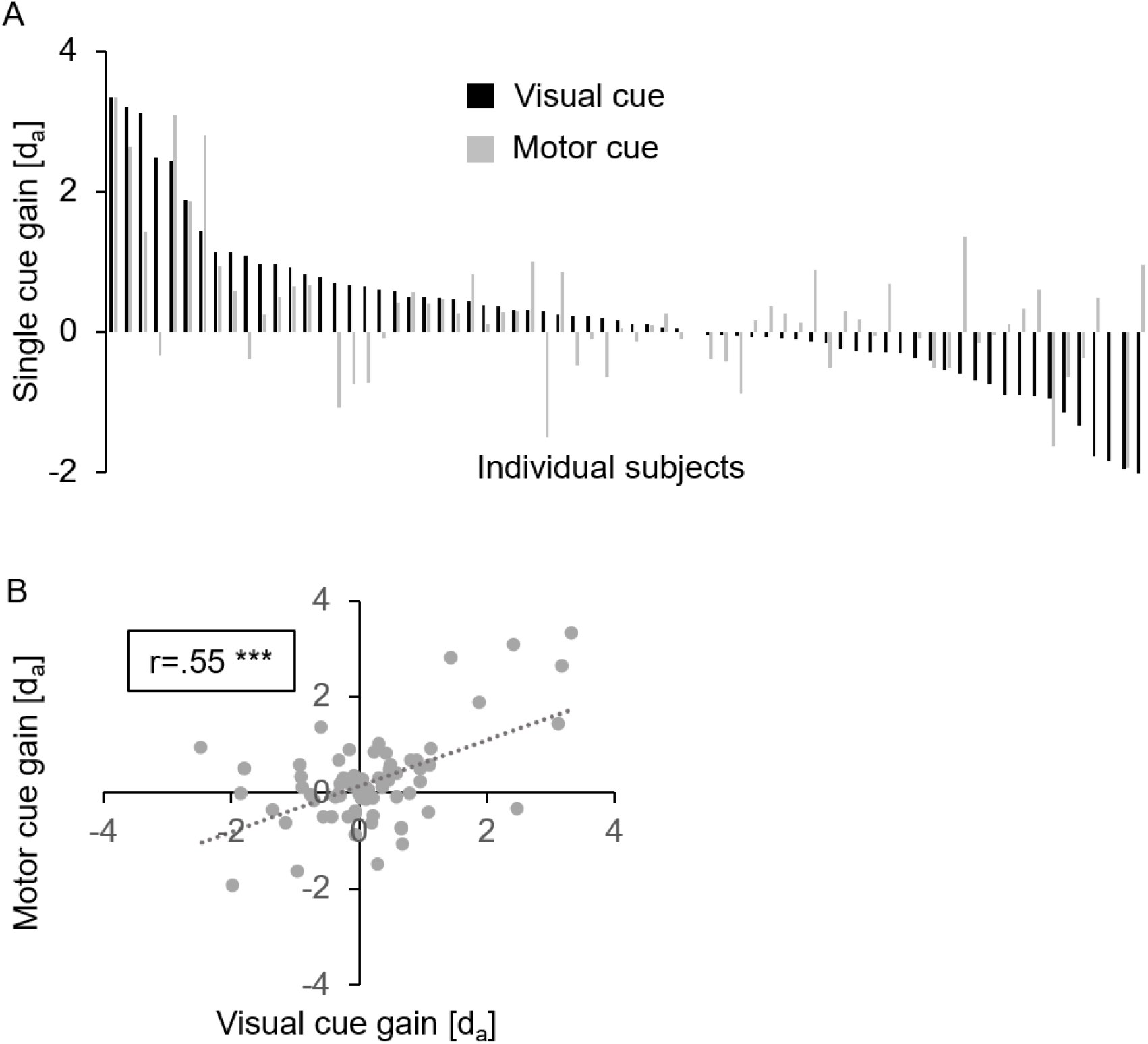
Memory gain in individual participants. A) Single cue gain calculated by subtracting the d_a_ values of the no cue condition from the d_a_ values of the single cue conditions: visual (black bars) or motor (grey bars). Each pair of gray and black bars represents an individual participant. Participants are sorted according to their visual gain (visual d_a_ minus no cue d_a_), starting from the participant who had the highest positive visual gain (on the left) to the participant who had the lowest negative visual gain (on the right). B) Scatter plot representing the visual cue gain (visual d_a_ minus no cue d_a_) relative to the motor cue gain (motor d_a_ minus no cue d_a_) in individual participants, and their fitted regression line (dashed line). *** p < .001.

### Good vs. Poor Performers

Next, we investigated whether the ability to use eye-related cues is linked to memory abilities. This was done by comparing the gain from cues between good and poor performers. Participants were divided into two groups based on median split on the average d_a_ value across all four conditions (median d_a_: 1.69). When examining each of the two groups separately, we found a significant correlation between visual cue gain and motor cue gain, in both groups (good performers: Pearson’s r = 0.54, p < 0.001; poor performers: Pearson’s r=0.58, p<0.001; **Fig. 4**). These results demonstrate that in both groups, some participants showed enhanced performance in the presence of the single cues, while others showed a decline compared to the no-cues condition. Thus, it seems that the ability to gain from the cues is not associated with memory ability.

**Figure 4.**
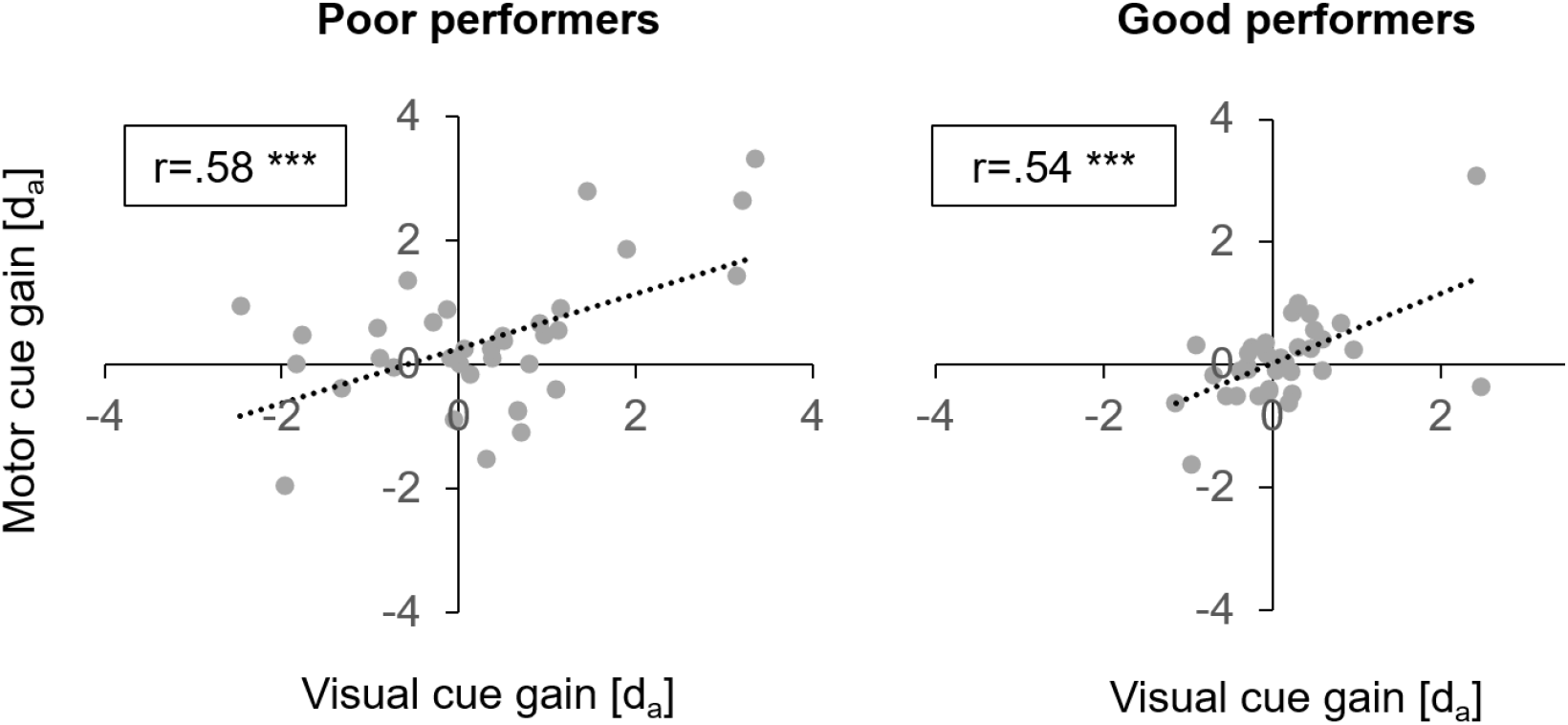
Memory gain in good vs. poor performers. Participants were divided into two groups, poor and good performance, according to a median split of the average d_a_ score of all four conditions. The scatter plot represents the visual cue gain (visual d_a_ minus no cue d_a_) relative to the motor cue gain (motor d_a_ minus no cue d_a_) in individual participants of both groups, and their fitted regression lines (dashed line). *** p < .001.

## Discussion

This study examined the role of the visual and the motor components of eye movements in memory retrieval. The findings show that both the visual and the motor components of eye movements contributed similarly to memory performance: no difference was found in the contribution of a single visual cue, a single motor cue, or the combination of both cues. Furthermore, the ability to gain from both types of eye movement-related cues varied substantially among individuals. Specifically, we found that individuals who gained from one cue tended to gain also from the other, while others gained from none or even experienced performance cost when cues were available. These findings provide first-time evidence for the roles of the motor and the visual components of eye movements as retrieval cues, and suggest that both components contribute to memory retrieval to a similar extent and that their relative contributions vary among individuals.

The findings support our hypothesis that eye movements act as memory retrieval cues because they lead to a match between encoding and retrieval in both the visual input and the motor activity. An alternative hypothesis was proposed by Richardson and Spivey^12^, who suggested that memory retrieval activates spatial pointers to the coordinates of the retrieved object. The activation of these pointers results in an eye movement toward the corresponding coordinates, aiming at collecting more data on the memorized item. This hypothesis emphasizes the importance of the visual, rather than the motor, aspect of eye movements as a memory retrieval cue. Our findings, on the other hand, highlight the key role of both of these aspects of eye movements in memory retrieval. Furthermore, by showing that externally-initiated eye movements enhance memory retrieval we show that eye movements can serve as an effective retrieval cue, at least during recognition of previously learned words.

The findings show that adding a visual eye movement-related cue over a motor cue or adding a motor cue over a visual cue did not enhance memory retrieval relative to a single cue. This finding is consistent with earlier studies, which investigated other types of retrieval cues, unrelated to eye movements. Evidence showed that when retrieval cues are presented sequentially, their effect was found to be aggregated^27–29^. In contrast, when two retrieval cues are presented simultaneously, no improvement in performance relative to a single cue is observed^30^. Indeed, in our study, the visual and motor eye-movement related cues in the VM condition represented two aspects of the same behavior and therefore, by definition, appeared simultaneously. It thus seems that under the simultaneous presentation in the VM condition, the two cues act as a single unified retrieval cue and hence lead to no benefit above and beyond any other single cue (V alone or M alone). Following Tulving and Osler^31^, this could indicate that when visual and motor information is available, eye movements constitute a single unified retrieval cue with integrated visual and motor components.

The findings further show substantial individual differences in the way people use eye related cues. This is consistent with previous studies that showed that individuals vary in the way they store and retrieve memories, and in the way they use memory retrieval cues^32–34^. It was found that people vary in their preferred learning style^35,36^, and that using this learning style leads to faster learning and better recall^37,38^. A few studies have suggested that individuals can be classified according to their preferred learning style as visual learners, auditory learners, motor learners and more^36,39^. Considering the visual and the motor components of eye movements, it could have been hypothesized that some individuals would rely more on the visual component, while others would rely more on the motor component. However, our findings showed a correlation between the gain from motor and visual cues, indicating that individuals tended to either benefit from both aspects of eye movements, or not benefit from any of them. This again supports the hypothesis that eye movement-related cues act as a single unified retrieval cue, which some individuals benefit from and some do not, as is the case for other types of retrieval cues.

A previous study found that there are substantial individual differences in the tendency to move the eyes during memory retrieval; some participants performed many saccades during retrieval while others hardly moved their eyes. Furthermore, evidence shows that this individual tendency to move one’s eyes during retrieval is not associated with enhanced memory performance ^32^. Unlike the previous study, the present study is not suitable for examining individual differences in the natural tendency to move the eyes because participants were asked to move their eyes in every trial. However, it could be hypothesized that the participants who gained less from eye movement related cues were the ones that naturally tend to make fewer eye movements. These participants may have developed other retrieval strategies and, perhaps, rely on different cues.

Counterintuitively, our results also reveal that some of the participants showed a consistent decline in performance following either a visual or a motor cue, relative to the no-cue condition. This finding seems to contradict the assumption that cues improve memory retrieval (when they are effective) or do not affect it (when they are ineffective). Why did the eye movement-related cues have negative effects on performance in some of the participants? The could answer be related to the binary nature of the cues in this design: there were only two options for the location of the presentation (top or bottom) and two for the direction of saccades (left-to-right or right-to-left). Therefore, the conditions in which there was no cue presented, there was, in fact, an *incongruent* cue: in the motor-cue condition, participants not only received a congruent motor cue but they also received incongruent visual information (top vs. bottom of the screen); in the visual-cue condition, participants not only received a congruent visual cue but they also received incongruent motor information (left vs. right saccades). Finding cost in these two conditions relative to the no-cue condition, in some participants, indicates that for these participants, performance was higher with two incongruent cues than with one congruent and one incongruent cue. In other words, more information led to lower memory performance.

This is not the first time in which findings showed that more information leads to less memory. Studies of the *partial list cueing effect* show that when encoding of items are presented as a list, providing participants with additional practice on a subset of this list, impairs memory performance for the rest of the items^40^. This was demonstrated in a variety of encoding conditions and types of tests, including in recognition tasks^41^. Research on the partial cueing effect suggests that the involved processes include inhibition for the unpracticed items^42^. Perhaps, in the present study, partial recreation of cues related to eye movements, while emphasizing the available cue, inhibited information that related to the missing cue. This could have resulted, in a minority of the individuals, in worse performance compared to the no cue condition. Finally, it is important to note that this counterintuitive cost from congruent cues, was found only in a minority of the participants. On average, participants performed better in the presence of single cues (motor or visual), indicating that most participants gained from eye movement-related cues.

## Conclusion

Understanding how we learn and remember and finding ways to enhance memory are some of the most important goals of cognitive research. This study revealed that eye movements, our most common and frequent behavior, play an important role in supporting memory. Furthermore, the present findings demonstrate how critical it is to consider individual differences in the use of memory retrieval cues: what helps one person may not necessarily help the other. Specifically, we found that some individuals tend to gain much from congruent eye movements during encoding and retrieval, while others do not gain at all or even suffer a cost. Recognizing this could advance research one step closer towards building personally-adjusted learning environments, that could help people improve learning and memory performance.

## Methods

### Participants

A total of 70 individuals participated in the experiment (53 females; ages 20-34, mean = 24.27, SD = 3.24). Sample size was chosen based on a power analysis using Gpower 3.1^43^, with an estimation of a medium effect size (Cohen’s d=0.5), with significance level of 0.05 and power of 0.8. Power estimation indicated a required sample size of N=64. Two additional participants were excluded from analysis because they did not provide sufficient data (see below). Participants were recruited through an online recruitment system and received course credit or payment for their participation. All participants had normal or corrected-to-normal vision and were native Hebrew speakers. Participants signed an informed consent prior to the experiment. The experiment was approved by the ethics committee of Tel Aviv University and of the School of Psychological Sciences.

### Procedure

Participants were provided with both written and verbal instructions regarding the experimental procedure and the eye-tracking calibration. The experiment included four blocks, each started with an encoding phase, followed by a break and then a recognition phase (**Fig. 4a**).

#### Encoding phase

56 encoding trials began with a black fixation cross (0.5°) appearing on one of the four corners of the screen (top-right, top-left, bottom-right or bottom-left) for 500ms. Next, a fixation cross disappeared and immediately reappeared at a new location, shifted 5.33° to the left or right of the first, on the horizontal axis. After 500ms, the fixation cross was replaced by a word (font Arial, size 0.25°X0.9°-1.2°) that appeared at the horizontal center of the screen at the same vertical location as the two crosses and remained there for 1000ms. The words were randomly chosen out a pre-defined dataset. The resulting experience of observers was of a fixation cross “jumping” twice on the horizontal axis, and then being replaced by the target word. Participants were asked to follow the fixation cross by performing two goal-directed saccades and to fixate and memorize the target word once it appears. They were informed that the experiment will advance to the next trial only after they fixate the stimulus on the screen. A break in fixation was defined as a 1^0^ distance from the stimuli, and when this happened the time for fixation was reset. Trials differed (in equal probability and at a pseudo-random order) by the direction of the saccades (from left to right or vice versa) and by the vertical position of the presentation (top/bottom). The encoding phase was followed by the break, in which participants were asked to fixate a red cross (1.3°) presented at the center of the screen for 10 seconds, and to prepare for the retrieval phase.

#### Retrieval phase

Retrieval trials were identical to those of encoding, with the exception that half the words had previously been presented at encoding (*target words*; 56 trials) and half had not (*lures*; 56 trials). Unlike in the encoding phase, the target word did not disappear after 1000ms, but remained on screen until the participant provided a response. For each word, participants were asked to determine whether it has been presented at encoding (‘old’) or not (‘new’), and to rate their confidence level in their decision on a scale from 1 to 3: 1 = low-confidence, 2 = moderate-confidence and 3 = high-confidence. Response was provided by pressing one of six keys: three keys on the right indicated the three confidence level ratings for “old” items, and three keys on the left indicated the three confidence level ratings for “new” items.

Retrieval trials belonged to one of the four cue conditions, as is depicted in **Fig. 4b**. The conditions are defined by the correspondence between their presentation in the encoding and the retrieval phases: a) **Visual (V) condition**: when the vertical position of stimulus presentation (top/bottom) was the same as in the corresponding encoding trials, but the direction of saccades (left/right) was opposite that of the corresponding encoding trials. b) **Motor (M) condition**: the direction of saccades was the same as in the corresponding encoding trials but the vertical position was opposite. c) **Visual-motor (VM) condition**: both the vertical position and the direction of saccades were the same as in the corresponding encoding trials. d) **No-cues condition**: both the saccades and the vertical position of the presentation were in the opposite direction to the corresponding encoding trials.

The experiment consisted of four blocks, each including target trials of only one of the four conditions. The order of the blocks was counterbalanced across participants. At the end of each block, participants were presented with a message informing them on the percentage of trials which were answered correctly (either “new” or “old”) out of the total number of trials in the block (0-100%).

Each word appeared only once in the experiment, either as a lure or a target. Targets and lures were chosen randomly for each participant out of a set of 448 Hebrew bi-syllabic nouns.

Prior to the main experiment, participants completed a training session that included a short version of a single block (with 16 targets), to ensure that they understand the task. Each block was about 12 minutes long. Participants were given the opportunity to take short breaks between blocks, if needed.

### Apparatus and Eye tracking

Participants were seated in a sound-attenuated room at a distance of 100 cm from a 24 inch ASUS VG248QE LCD screen, with their head rested on a chin-rest.

Eye-movements were monitored binocularly using a remote infrared video-oculographic system (EyeLink 1000 Plus, SR Research Ltd, Ontario, Canada), with a sampling-rate of 1000 Hz, spatial resolution <0.01° and average accurate of 0.25°-0.5° average accuracy when using a head-rest, as reported by the manufacturer.

At the beginning of each experiment, participants completed a 9-point eye tracking calibration procedure performed with the same illumination conditions as the main experiment. Calibration was accepted when validation confirmed that fixation errors were of no more than 0.2°. Calibration task was repeated as needed later in the experiment.

### Analysis

The analysis was based on signal-detection theory, SDT^25^. The classic index of d’ assumes equal variance of the lure and target distributions, which has been shown to not be a valid assumption for recognition memory^44^. Critically, it has been shown that when indices of sensitivity are used under false assumption, response bias effects may be wrongly interpreted as those of sensitivity^45^. We thus used a related SDT index, d_a_. Like d’, d_a_ provides an estimate of the distance between the means of the lure and target distribution in units of their shared standard deviation. However, d_a_ does not assume equality of variance. Rather, it depends on estimating the inequality of variance with a measure, S, and incorporating S into the sensitivity index (see next paragraph). S is an estimate of the lure-to-target distribution standard deviations. Importantly, for d’, only binary responses (old/new) are needed. In contrast, d_a_ relies on confidence ratings reported by participants in each trial (in our case, a total of six levels - three per category), from which it is possible to estimate S. In all, we calculated for each participant four d_a_ values, one for each of the four conditions^26^.

The measure S is equal to the slope of the z-transformation of the receiver operating characteristic (zROC). To obtain this, we computed hit rates (“H”, in trials with target words) and false-alarms rates (“FA”, in trials with lures) corresponding to five levels of response bias. For the most conservative level, only responses made at the highest level (confidence level 6, CL6) were regarded as ‘old’. For each participant, we counted the number of targets and lures made at CL6, to obtain the first operating point on the ROC curve. For a slightly less conservative response bias, all targets and lures made at either CL5 or CL6 were regarded as ‘old’, yielding pair of H and FA. We continued this way until the most liberal operating point, regarding as ‘old’ all CLs from 2-6. Wet thus obtained a total of five pairs of H-FA for each of the four experimental conditions, for each participant.

It has been shown that if the mnemonic signal has a Gaussian distribution, then displaying the ROC curve in Z space yields a linear curve^46^. Critically, the slope of zROC curve, S, provides a good estimate of the ratio of the lure-to-target distribution standard deviations^47^. The estimate S represents the average of the slopes of the zROC curves of the four conditions, per participant. Since calculating the slope relies on having minimal variability of the confidence ratings, in cases in which participants provided only one or two levels of confidence ratings the slope could not be estimated. In these cases, S was calculated as the mean of those conditions for which the slope could be estimated. In cases (N = 2) where all four slopes were incalculable, the participant was excluded from analysis.

Using the value S and the original hits (H) and false alarm (FA) rates as set by the ‘old’ and ‘new’ responses across their respective levels of confidence, d_a_ was calculated as follows:

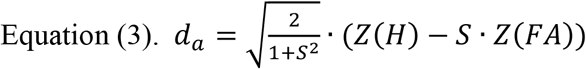

We calculated the d_a_ values and slopes for each participant and condition. Because all H and FA rates are calculated using the Gaussian distribution, calculations of Z scores of 100% and 0% provide NaN-values. To solve this, we followed the proposition made by Hautus^48^: for blocks with 100% hits, hit rate was calculated as (targets+0.5)/(targets+1) (in our experiment, resulting in hit rates of 0.9912). For blocks with 0% of false alarms, calculation was the opposite: (0.5)/(lures+1) (in our experiment, resulting in false alarm rates of 0.0088).

## Acknowledgements

This research was funded by the Israel Science Foundation, grant 1960/19 to S.Y-G and by the Ariane de Rothschild Women Doctoral scholarship to K.T.

## Authors contribution

SYG, KT and YGG designed the research; KT performed the research; KT and YGG analyzed data; SYG, KT and YGG wrote the paper.

## Data availability statement

The data that support the findings of this study are available from the corresponding author, KT, upon reasonable request.

## References

1. Abeles, D. & Yuval-Greenberg, S. Just look away: Gaze aversions as an overt attentional disengagement mechanism. Cognitions 168, 99–109 (2017).

2. Brandt, S. A. & Stark, L. W. Spontaneous Eye Movements During Visual Imagery Reflect the Content of the Visual Scene. J. Cogn. Neurosci. 9, 27–38 (1997).

3. Ferreira, F., Apel, J. & Henderson, J. M. Taking a new look at looking at nothing. Trends Cogn. Sci. 12, 405–410 (2008).

4. Renkewitz, F. & Jahn, G. Looking at Nothing Indicates Memory Search in Multiattribute Decision Making. Proc. Annu. Meet. Cogn. Sci. Soc. 32, 260–265 (2010).

5. Spivey, M. J. & Geng, J. J. Oculomotor mechanisms activated by imagery and memory: eye movements to absent objects. Psychol. Res. 65, 235–241 (2001).

6. Altmann, G. T. M. Language-mediated eye movements in the absence of a visual world: the ‘blank screen paradigm’. Cognition 93, B79–B87 (2004).

7. Henderson, J. M. & Castelhano, M. S. Eye movements and visual memory for scenes. in Cognitive processes in eye guidance (ed. Underwood, G.) 213–235 (Oxford University Press, 2005).

8. Hoover, M. A. & Richardson, D. C. When facts go down the rabbit hole: Contrasting features and objecthood as indexes to memory. Cognition 108, 533–542 (2008).

9. Irwin, D. E. Memory for Position and Identity Across Eye Movements. 18, 307–317 (1992).

10. Johansson, R., Holsanova, J. & Holmqvist, K. Pictures and Spoken Descriptions Elicit Similar Eye Movements During Mental Imagery, Both in Light and in Complete Darkness. Cogn. Sci. 27 (2006).

11. Zelinsky, G. J. & Loschky, L. C. Eye movements serialize memory for objects in scenes. Percept. Psychophys. 67, 676–690 (2005).

12. Richardson, D. C. & Spivey, M. J. Representation, space and Hollywood Squares: looking at things that aren’t there anymore. Cognition 76, 269–295 (2000).

13. Brockmole, J. R. & Irwin, D. E. Eye movements and the integration of visual memory and visual perception. Percept. Psychophys. 67, 495–512 (2005).

14. Melcher, D. & Kowler, E. Visual scene memory and the guidance of saccadic eye movements. Vision Res. 15 (2001).

15. Laeng, B. & Teodorescu, D.-S. Eye scanpaths during visual imagery reenact those of perception of the same visual scene. Cogn. Sci. 26, 207–231 (2002).

16. Johansson, R. & Johansson, M. Look Here, Eye Movements Play a Functional Role in Memory Retrieval. Psychol. Sci. 25, 236–242 (2014).

17. Bradley, M. M., Cuthbert, B. N. & Lang, P. J. Perceptually driven movements as contextual retrieval cues. Bull. Psychon. Soc. 26, 541–543 (1988).

18. Bochynska, A. & Laeng, B. Tracking down the path of memory: eye scanpaths facilitate retrieval of visuospatial information. Cogn. Process. 16, 159–163 (2015).

19. Scholz, A., Mehlhorn, K. & Krems, J. F. Listen up, eye movements play a role in verbal memory retrieval. Psychol. Res. 80, 149–158 (2016).

20. Tulving, E. Elements of Episodic Memory. (Oxford University Press, 1983).

21. Tulving, E. & Thomson, D. M. Encoding specificity and retrieval processes in episodic memory. Psychol. Rev. 80, 352–373 (1973).

22. Spivey, M. J. The continuity of mind. (Oxford University Press., 2008).

23. Pylyshyn, Z. The role of location indexes in spatial perception: A sketch of the FINST spatial-index model. Cognition 32, 65–97 (1989).

24. Richardson, D. C. & Kirkham, N. Z. Multimodal Events and Moving Locations: Eye Movements of Adults and 6-Month-Olds Reveal Dynamic Spatial Indexing. J. Exp. Psychol. Gen. 133, 46–62 (2004).

25. Tanner, W. P. & Swets, J. A. A decision-making theory of visual detection. Psychol. Rev. 61, 9 (1954).

26. Didi-Barnea, C., Peremen, Z. & Goshen-Gottstein, Y. The unitary zROC slope in amnesics does not reflect the absence of recollection: critical simulations in healthy participants of the zROC slope. Neuropsychologia 90, 94–109 (2016).

27. Massaro, D. W., Weldon, M. S. & Kitzis, S. N. Integration of Orthographic and Semantic Information in Memory Retrieval. J. Exp. Psychol. Learn. Mem. Cogn. 17, 277–287 (1991).

28. McLeod, P. D., Williams, C. E. & Broadbent, D. E. Free Recall with Assistance from One and from Two Retrieval Cues. Br. J. Psychol. 62, 59–65 (1971).

29. Nelson, D. L., Fisher, S. L. & Akirmak, U. How implicitly activated and explicitly acquired knowledge contribute to the effectiveness of retrieval cues. Mem. Cognit. 35, 1892–1904 (2007).

30. Tulving, E. & Osler, S. Effectiveness of retrieval cues in memory for words. J. Exp. Psychol. 77, 593–601 (1968).

31. Tulving, E. & Osler, S. Effectiveness of retrieval cues in memory for words. J. Exp. Psychol. 77, 593–601 (1968).

32. Kinjo, H., Fooken, J. & Spering, M. Do eye movements enhance visual memory retrieval? Vision Res. 176, 80–90 (2020).

33. Sahakyan, L., Abushanab, B., Smith, J. R. & Gray, K. J. Individual differences in contextual storage: Evidence from the list-strength effect. J. Exp. Psychol. Learn. Mem. Cogn. 40, 873–881 (2014).

34. Unsworth, N. Individual differences in long-term memory. Psychol. Bull. 145, 79–139 (2019).

35. Lodge, J. M., Hansen, L. & Cottrell, D. Modality preference and learning style theories: rethinking the role of sensory modality in learning. Learn. Res. Pract. 2, 4–17 (2016).

36. Magoon, R. A. & Raper, C. C. The effect of sensory modalities on learning. Contemp. Educ. Psychol. 2, 55–65 (1977).

37. Korenman, L. M. & Peynircioglu, Z. F. Individual Differences in Learning and Remembering Music: Auditory versus Visual Presentation. J. Res. Music Educ. 55, 48–64 (2007).

38. Moura, H. Learning Styles: From History to Future Reasearch Implications for Distance Learning. Rev. Bras. Aprendiz. Aberta E Distância 6, (2008).

39. Barbe, W. B. & Milone Jr, M. N. What We Know About Modality Strengths. Educ. Leadersh. 38, 378–380 (1981).

40. Slamecka, N. J. An examination of trace storage in free recall. J. Exp. Psychol. 76, 504–513 (1968).

41. Todres, A. K. & Watkins, M. J. A Part-Set Cuing Effect in Recognition Memory. J. Exp. Psychol. 7, 91–99 (1981).

42. Aslan, A., Bäuml, K.-H. & Grundgeiger, T. The role of inhibitory processes in part-list cuing. J. Exp. Psychol. Learn. Mem. Cogn. 33, 335–341 (2007).

43. Faul, F., Erdfelder, E., Lang, A.-G. & Buchner, A. G*Power 3: A flexible statistical power analysis program for the social, behavioral, and biomedical sciences. Behav. Res. Methods 39, 175–191 (2007).

44. Mickes, L., Wixted, J. T. & Wais, P. E. A direct test of the unequal-variance signal detection model of recognition memory. Psychon. Bull. Rev. 14, 858–86 (2007).

45. Rotello, C. M., Heit, E. & Dubé, C. When more data steer us wrong: replications with the wrong dependent measure perpetuate erroneous conclusions. Psychon. Bull. Rev. 22, 944–954 (2015).

46. Yonelinas, A. P. Recognition memory ROCs for item and associative information: The contribution of recollection and familiarity. Mem. Cognit. 25, 747–763 (1997).

47. Hautus, M. J., Macmillan, N. A. & Rotello, C. B. Toward a complete decision model of item and source recognition. Psychon. Bull. Rev. 15, 889–905 (2008).

48. Hautus, M. J. Corrections for extreme proportions and their biasing effects on estimated values ofd’. Behav. Res. Methods Instrum. Comput. 27, 46–51 (1995).

